# Single human fingertip mechanoreceptive afferents simultaneously encode multidimensional aspects of touch

**DOI:** 10.1101/2025.10.28.685066

**Authors:** Victoria Ashley Lang, Helena Backlund Wasling, Rochelle Ackerley, Johan Wessberg

**Affiliations:** University of Gothenburg, Inst. of Neuroscience and Physiology, Dept. of Physiology; Aix Marseille Univ, CNRS, CRPN (Centre de Recherche en Psychologie et Neurosciences – UMR 7077), Marseille, France

**Keywords:** human, tactile, microneurography, low-threshold mechanoreceptor, skin, computational modeling, regression analysis

## Abstract

Touch using the hands is essential for recognizing surface features and manipulating objects, where different aspects are encoded by four main types of low-threshold mechanoreceptors (LTMs) in glabrous skin. Although factors such as movement, vibration, and pressure are often studied individually, touch requires their integration to produce complex sensations. We investigated these processes jointly by recording single-unit activity via microneurography from human LTM afferents in the median nerve as periodic gratings slid across their receptive fields, varying the normal force and sliding velocity per trial. Mixed-effects models revealed that fast-adapting type 1 (FA-1) afferent firing was influenced by all three parameters—force, velocity, and spatial period. Slowly-adapting type 1 (SA-1) afferent firing was primarily driven by force, and to a lesser extent, velocity. These findings support the view that FA-1 afferents encode stimulus intensity in an approximately linear manner, while SA-1 afferents function mainly as force detectors, demonstrating that mechanoreceptive afferents provide a complementary and multidimensional representation of texture during sliding contact. This distributed encoding challenges the notion that LTMs are dedicated to single stimulus features, suggesting that tactile information is represented across LTM populations where each class contributes differently-weighted inputs to capture skin-object interactions.

## Introduction

Our hands are central to how we explore and interact with the world, enabling us to perceive objects and textures with remarkable precision. This mechanical contact occurs over a wide range, where the skin moves dynamically in three dimensions under various normal, tangential, and lateral forces, which is differentially encoded by classes of low-threshold mechanoreceptive (LTM) afferents in the skin. In typical tactile interactions, the finger pad often slides over surfaces of diverse textures, which have specific vibrational components, meaning we can readily perceive differences between silk and sandpaper, but also between silk and satin. Understanding how tactile information is encoded in the glabrous skin of the hand, especially at the finger pad, is therefore essential for explaining the neural basis of fine touch perception. Touch processing mechanisms have been extensively studied in animal models^1–4^, providing crucial insights into the neurophysiological basis of human tactile discrimination. However, this does not always directly translate and how human mechanoreceptive afferents encode dynamic stimuli is an active area of investigation, where the technique of microneurography in human peripheral nerves enables direct recordings from LTMs, bridging knowledge from animal research to humans^5–7^.

Studies using microneurography have identified four types of fast-conducting LTM afferents in human glabrous skin: fast-adapting type 1 (FA-1, Meissner), fast-adapting type 2 (FA-2, Pacinian), slowly-adapting type 1 (SA-1, Merkel), and slowly-adapting type 2 (SA-2, Ruffini)^8,9^. Each afferent type is selectively tuned to particular stimulus features and plays complementary roles in encoding tactile information^8,9^. Classic studies have described their receptive field properties and adaptation behaviors, but questions remain about how these afferents encode complex, dynamic touch. Foundational works in non-human primates have been instrumental in shaping our current understanding of texture encoding at the level of single afferents. Seminal studies^10,11^ examined how LTM afferents in macaques responded to spatial features of textured surfaces, demonstrating that SA-1 afferents convey fine spatial detail by producing spatially stable firing patterns that mirror the geometry of raised-dot and grating stimuli, whereas FA-1 afferents respond more transiently to motion and edges, emphasizing changes rather than static form. Later research extended these findings by showing that tactile afferents convey information not only through mean firing rates, but also through precise temporal spiking timing, capturing fine surface features and vibration-induced microstructure^12–14^. More recent work has shown that naturalistic textures elicit richly patterned responses in human and monkey afferent populations, and that neurons in the somatosensory cortex integrate these temporal codes to build higher-level representations of texture^15,16^. These studies have established fundamental principles: the role of temporal precision, population coding, and spike timing in fine texture discrimination and slip detection^17,18^. Yet, it remains unclear to what extent these coding principles generalize to human touch, given differences in afferent distributions and perceptual thresholds^19,20^.

In the last decade, computational models have been formulated to address these gaps^15,18,21^. However, widely used models, such as TouchSim^18^ that is based on monkey recordings, do not fully capture the workings of the human system, limiting their applicability to human tactile encoding. Clarifying whether single human mechanoreceptive afferents encode temporal structure through precise spike timing or primarily via firing rate modulation remains central to explaining fine texture discrimination. Moreover, how LTM afferents integrate dynamic factors during sliding contact—such as normal force, velocity, and spatial period—has been little explored. In the present study, we hypothesized that individual LTM afferents show varying sensitivity to multiple dynamic features and are not exclusively tuned to single stimulus parameters. Thus, we recorded from single LTM afferent units in the human fingertip as periodic gratings were passed across their receptive fields, varying their spatial period, normal force, and sliding velocity, aiming to determine the relative contributions of each parameter to mechanoreceptor activity.

## Results

A total of 27 single-unit LTM afferent units were recorded from the upper median nerve, while a mechatronic platform delivered periodic gratings with controlled normal force and sliding velocity (Fig. 1A). These recordings were made from 13 FA-1, 2 FA-2, 10 SA-1, and 2 SA-2 afferents (Fig. 1B). Due to fewer recordings from type 2 LTMs (SA-2 and FA-2), the analyses primarily focus on the more numerous type 1 (FA-1 and SA-1) afferents, which provided sufficient data for reliable statistical modeling.

**Figure 1.**
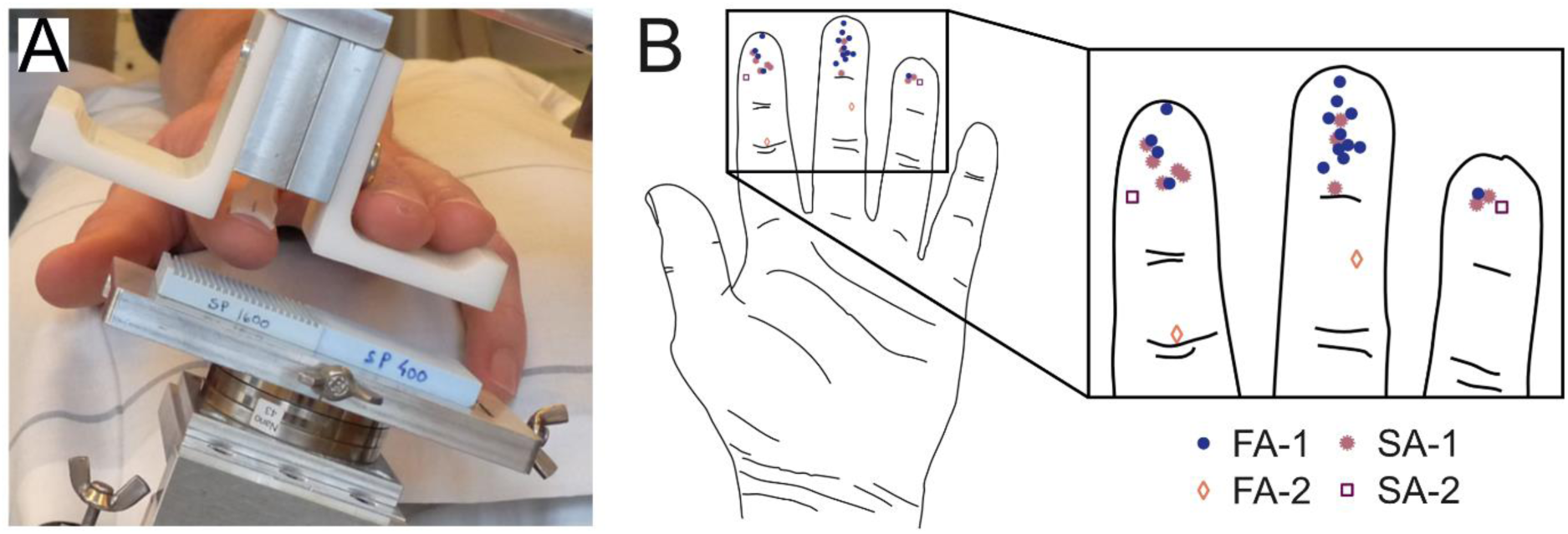
Setup and schematic of the low-threshold mechanoreceptive (LTM) afferents recorded from. (A) The mechatronic platform with a plate on which two periodic gratings are mounted, atop and perpendicular to the receptive field of an LTM located on the index finger pad, which is fixed to a holder. (B) Schematic of a hand with the 27 LTM afferent units that we recorded from on the upper volar surfaces of the three middle fingers, with a zoomed-in view on the right.

We obtained data over four different sliding velocities (5, 10, 20, 40 mm/s) and at four different normal forces (100, 200, 400, or 800 mN) using gratings of 17 different spatial periods. Figure 2 provides an example of a single run for the FA-1 unit N092 and SA-1 unit N144 both under 200 mN normal force, 20 mm/s sliding velocity, and 400 μm grating. The figure shows the onset and offset of tactile contact with a corresponding burst of spikes each time, where a stronger response was often seen for the initial touch onset and the activity was much higher for the FA-1. The middle of the figure illustrates the response to sliding of the grating across the receptive field of the FA-1 unit that was located on the tip of the index finger. A transient, high peak in FA-1 afferent firing corresponding to the onset of sliding was often observed that continued as regular firing while the grating was moved over the skin, showing a baseline firing rate of around 40 spikes/s, with brief peaks of activity throughout. Conversely, the activity from the SA-1, shown at the bottom of the figure, is far less intense, thus showing lower variability, and the initial peak was far less pronounced. These were typical modes of response for these two types of LTM.

**Figure 2.**
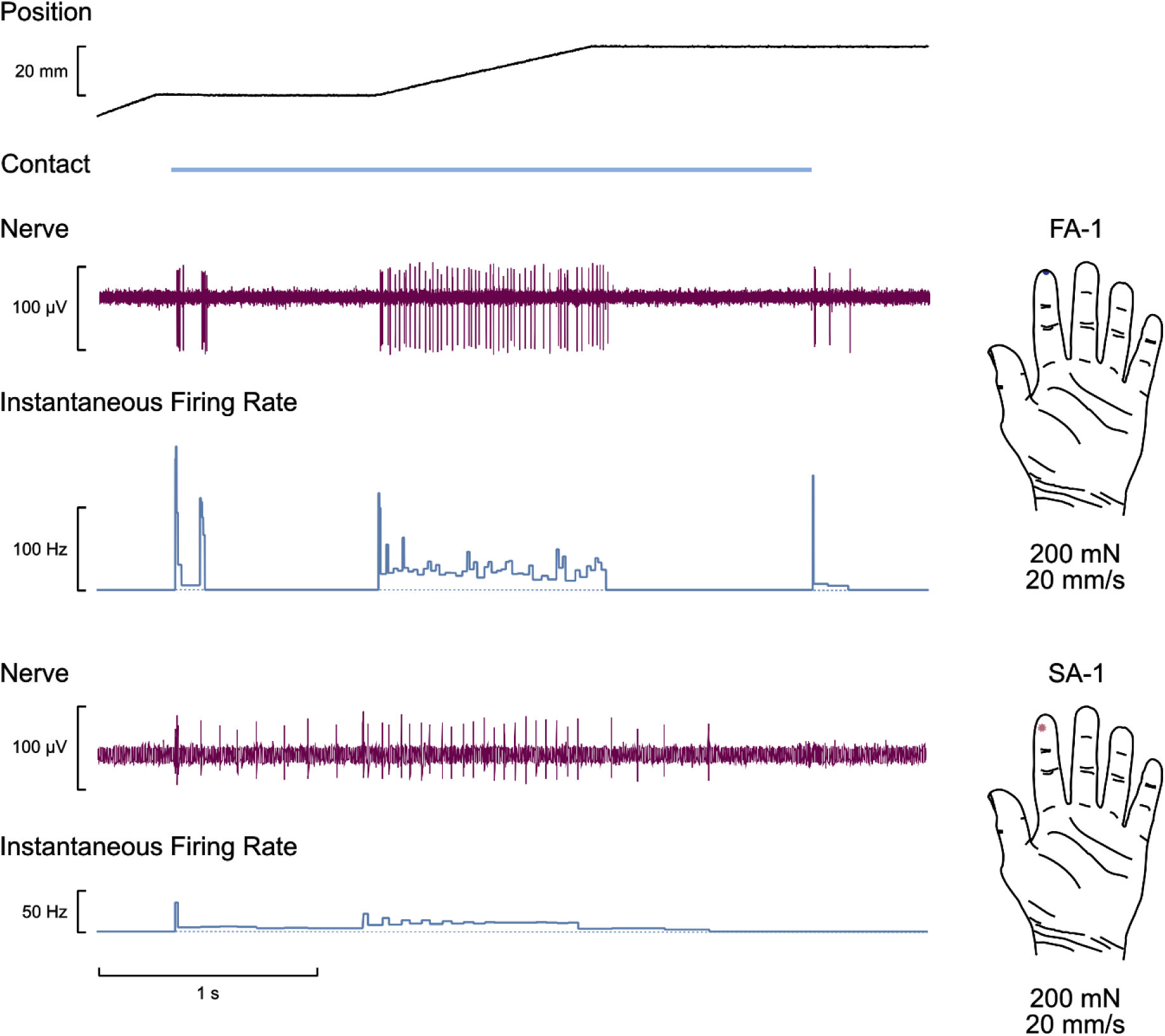
An example of a single run with the robotic platform for FA-1 unit N092 and SA-1 unit N144 located on the tip of the index finger under 200 mN normal force, 20 mm/s sliding velocity, and 400 μm grating. From top to bottom: position of grating, contact period with skin, microneurographic nerve recording, and instantaneous firing rate (IFR).

In comparing differences over spatial periods moved over the receptive field, Figure 3 illustrates examples of spike raster plots showing the variation in response patterns across an increase in spatial period for different FA-1 and SA-1 units. Figure 3A demonstrates the FA-1 unit’s spike activity, at a controlled 400 mN force and 10 mm/s sliding velocity, where an overall increase in firing rate can be seen, as the spatial period increased. However, the correspondence between the spatial period and the unitary response is not direct, as seen in the variability over the 12 repetitions for each of the three spatial periods. The response appears somewhat irregular and only partly reflects the grating’s periodic structure. Similarly, Figure 3B presents an SA-1 unit’s spike activity, at a controlled 800 mN force and 10 mm/s sliding velocity, where firing rate also increased as a function of spatial period. Here, the unit fired more regularly only for the coarser gratings of 1600 and 1920 µm spatial period, suggesting that larger ridge spacing more effectively engaged its receptive field.

**Figure 3.**
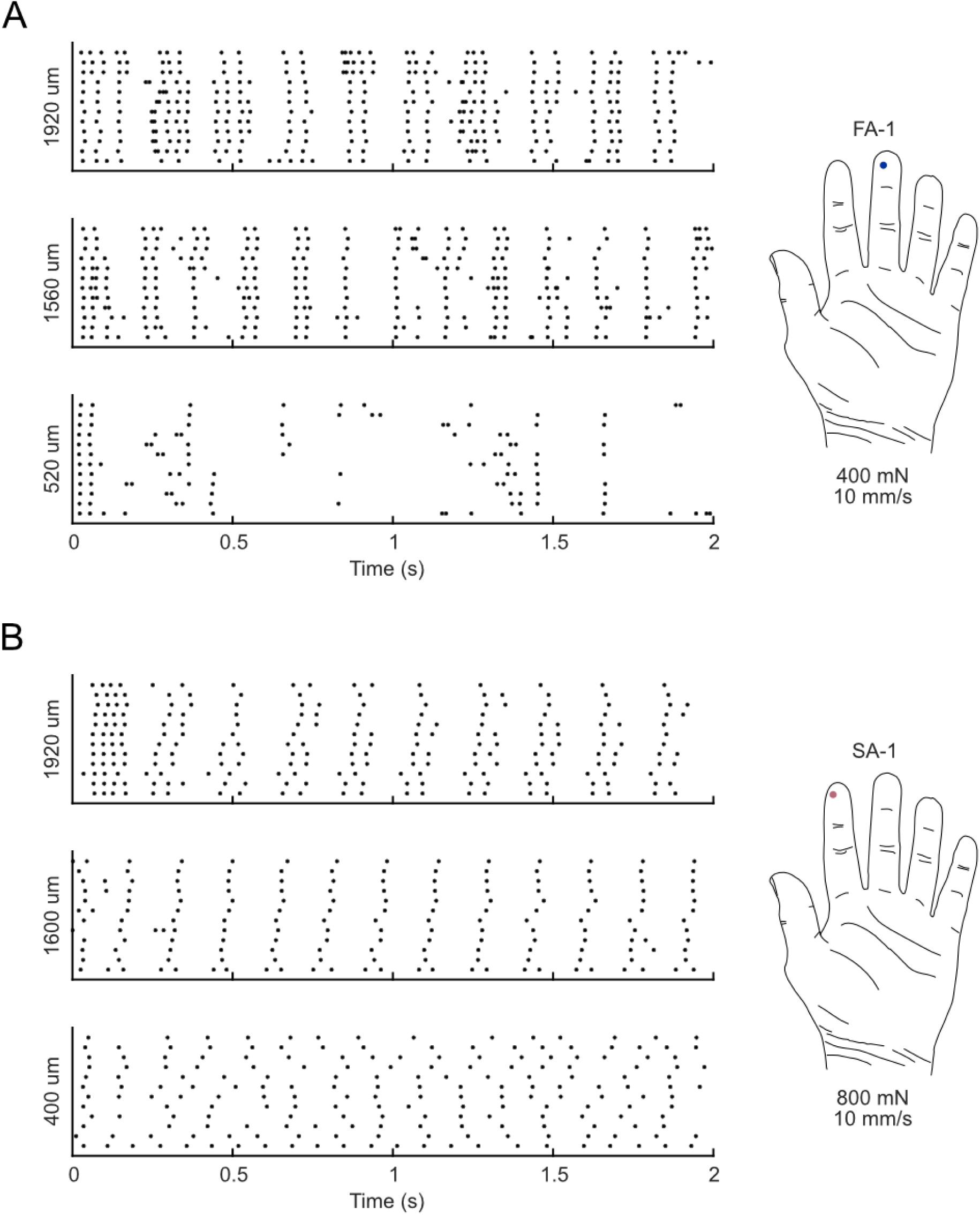
Differences in firing with spatial period. Spike raster plots for (A) FA-1 unit N111 subject to 400 mN normal force and 10 mm/s sliding velocity, and (B) SA-1 unit N191 subject to 800 mN normal force and 10 mm/s sliding velocity for 3 gratings of varying spatial period. Both units had 12 repetitions of each stimulus.

Figure 4 presents boxplots of median instantaneous firing rates (IFR) across four sliding velocities for different FA-1, SA-1, FA-2 and SA-2 units. The FA-1 shows a clear increase in median IFR with increasing sliding velocity and its increase with spatial period can also be seen (Fig. 4A), indicating its sensitivity to both dynamic and spatial features. The SA-1 unit (Fig. 4B) shows a similar trend, but with narrower firing rate ranges, consistent with its known role as a force-sensitive, slowly-adapting receptor. For the limited FA-2 and SA-2 data (Fig. 4C-D), no distinct trends could be established concerning the responses to increasing sliding velocity. However, it can be noted that there was a marked difference in IFR between fast– and slowly-adapting types, where the FA-1 and FA-2 reached firing frequencies of well over 100 spikes/s, whereas the SAs peaked around 20 spikes/s.

**Figure 4.**
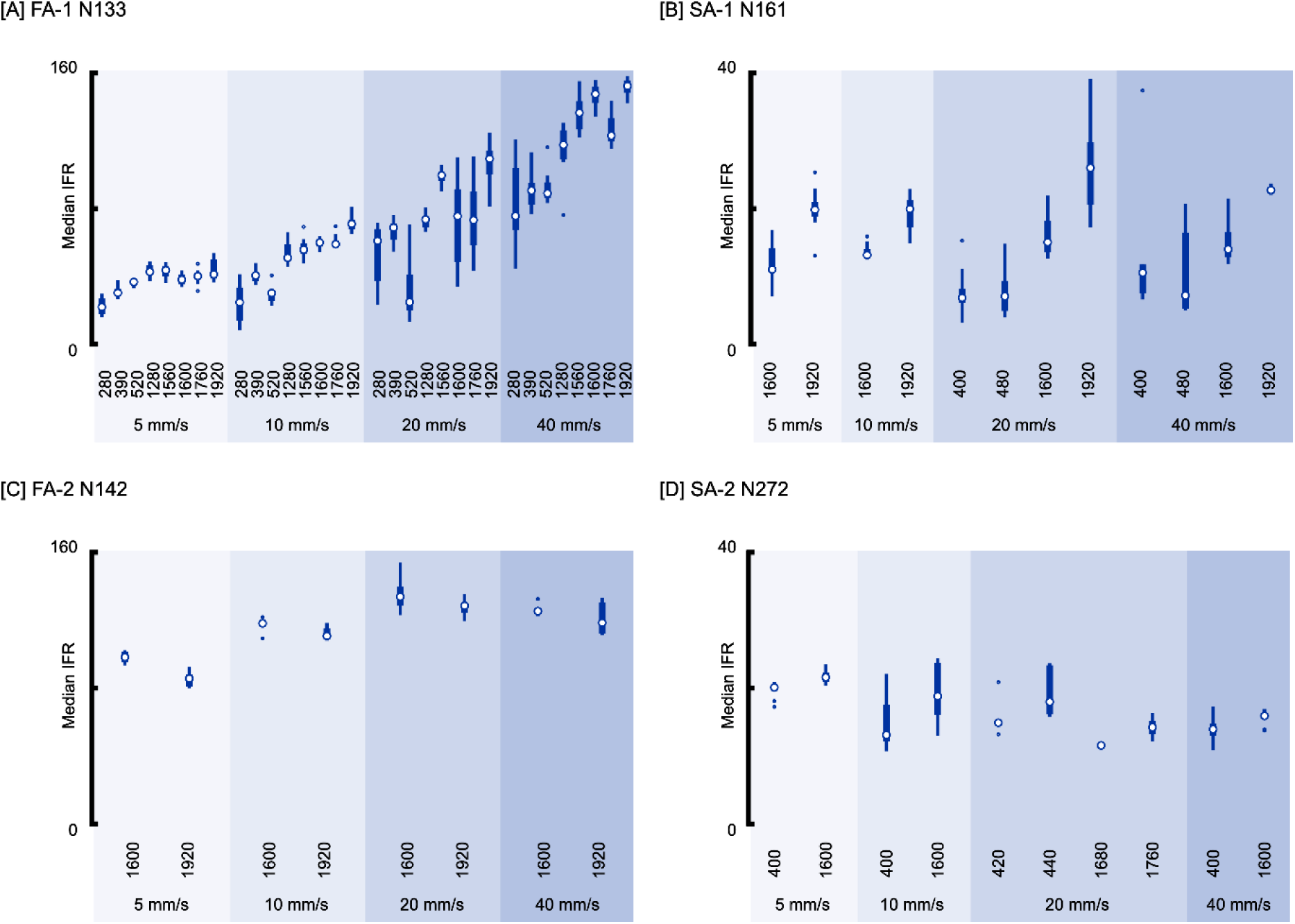
Differences in firing for sliding velocity, over different spatial periods. Boxplots show the median instantaneous firing rates (IFR) for (A) FA-1 unit N133 subject to 200 mN, (B) SA-1 unit N161 subject to 400 mN, (C) FA-2 unit N142 subject to 200 mN, and (D) SA-2 unit N272 subject to 400 mN. The median IFR appears to increase as a function of velocity and spatial period for these FA-1, SA-1, and SA-2 units. The lower and upper edges of the box demarcate the interquartile range, and the ‘bullseye’ indicates the median. Note that the median IFR scales are the same for the FA afferents and for the SA afferents.

An overview of median IFRs over the conditions is presented in Figure 5 for a different FA-1 unit, across two normal forces, four sliding velocities, and 8 spatial periods. Logarithmic curves were fit to each sliding velocity condition, highlighting the increase in firing rate with higher velocity and with coarser spatial period. Comparing the 200 mN and 400 mN force conditions (blue curves in Figure 5) shows the increase in median IFR with elevated force. The above curves are not illustrated for the other unit types, due to the difficulty in having a large number of recorded combinations in any one unit.

**Figure 5.**
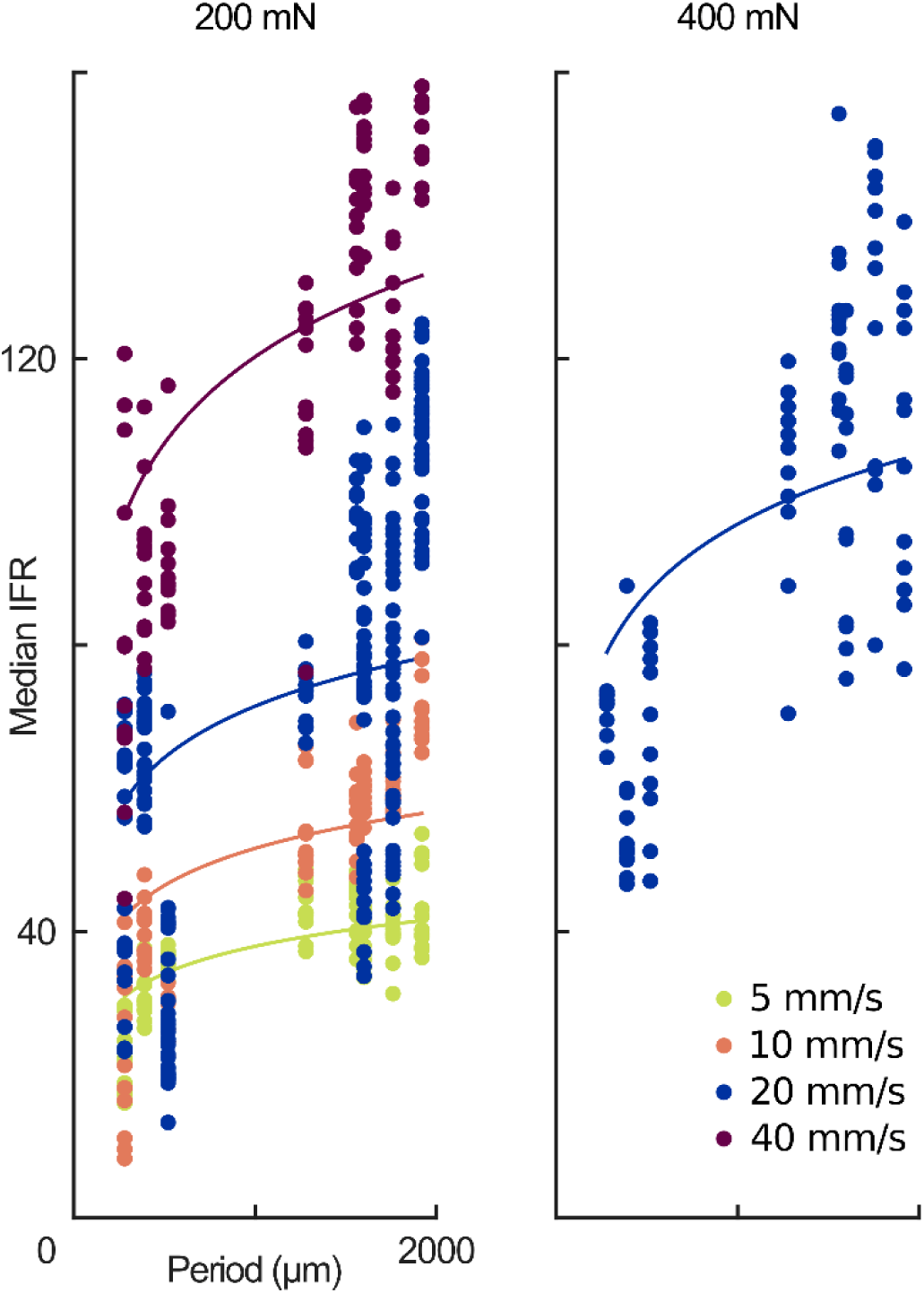
Differences in firing with sliding velocity, over different forces and spatial periods. Median instantaneous firing rates (IFR) overview for FA-1 unit N111 at applied normal forces of 200 and 400 mN. For each velocity condition, logarithmic curves detail the firing rate trend as a function of 8 gratings of different spatial periods. The median IFR generally increases as a function of all three stimulus parameters.

To understand better how these parameters combined to produce the different firing responses, we conducted analyses per afferent type, namely on the type 1 units where more trials had been collected. We assessed the similarity between the FA-1 afferent units (n = 13), performing single-level multivariate regressions for each FA-1 unit (Fig. 6A). These regressions confirmed that FA-1 units exhibit increased firing with the combined influence of the log-transformed stimulus parameters. This result supported the construction of mixed-effects models to evaluate how well each stimulus parameter predicts median IFR across units. Individual units were treated as the grouping factor, and the combination of stimulus parameters gave rise to three multilevel models using a single predictor, three models using two predictors, and one model utilizing all stimulus parameters. A comparison of the models suggests that FA-1 afferent firing is responsive to changes in all three stimulus parameters—force, velocity, and spatial period. That is, an increase in any of the three stimulus parameters results in an increase in the median IFR. Model comparisons (Figs. 6B-C) show that the full model, incorporating all three parameters, yielded the narrowest coefficient estimates, lowest AIC and highest R^2^ values, indicating that FA-1 afferents integrate force, velocity, and texture features together in an additive manner, in which we explore their contribution below.

**Figure 6.**
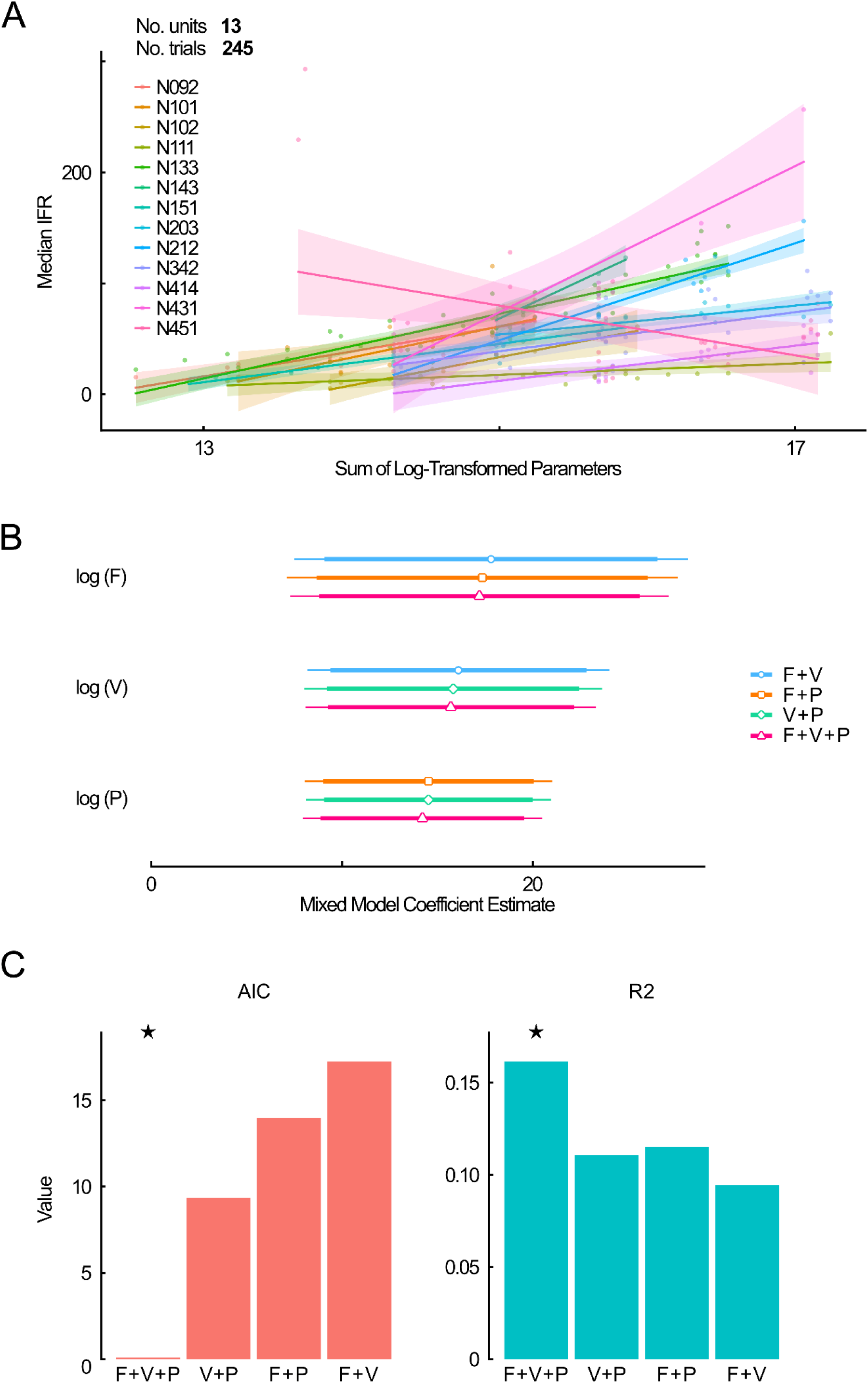
(A) Individual regression models for each FA-1 unit show the contribution of stimulus parameters—force, velocity, and spatial period—to the median IFR. Thirteen units formed 245 trials. (B) A comparison of predictor coefficients for mixed models shows that the complete model (pink) has the smallest coefficient uncertainty (narrowest confidence interval). (C) The complete model (marked by the stars) gives the lowest AIC and highest R-squared. The AIC values are computed as the difference in AIC to the starred model for better visualization.

We also assessed the firing at the group level of individual SA-1 afferents (Fig. 7). Although 10 SA-1s were included in the analysis, the single-level multivariate regressions appear less consistent than the FA-1s for a mixed-effects model development, especially in four SA-1s where there was higher variability and/or fewer parameters tested (see blue and green lines in Fig. 7A). However, on comparing the mixed-effects models, it appears that SA-1 firing is primarily driven by force and velocity, with spatial period showing negligible influence, as shown in Figure 7. In comparing the models (Figs. 7B-C), it is evident that the full model and the force + velocity model perform the best. The force + velocity model yields the lowest AIC, with the full model close by, whereas the highest R^2^ are from the full model, with the force + velocity model nearby. This indicates that SA-1 afferents mainly integrate force and velocity information, with spatial period influencing the firing to a lesser extent.

**Figure 7.**
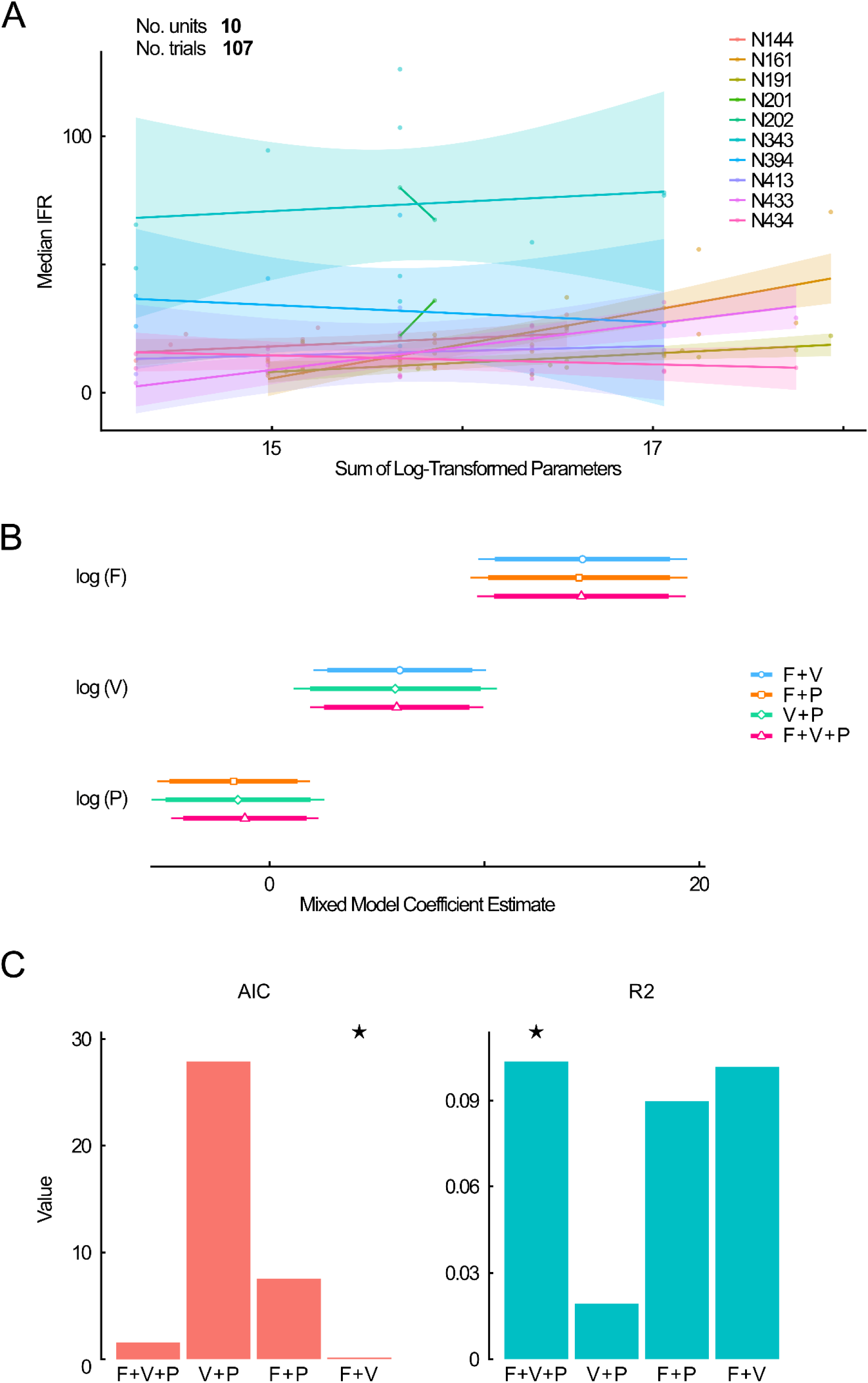
(A) Individual regression models for each SA-1 unit show the contribution of stimulus parameters—force, velocity, and spatial period—to the median IFR. Ten units formed 107 trials. (B) A comparison of predictor coefficients for mixed models showed that the two-predictor force-velocity model (blue) and the complete model (pink) have the smallest coefficient uncertainties (narrowest confidence interval). (C) The preferred models according to AIC and R^2^ are marked with a star. The AIC values are computed as the difference in AIC to the starred model for better visualization.

To validate the robustness of the FA-1 and SA-1 findings, we used bootstrapping analyses, performed with 1000 resamples. In line with the above results, the predictor coefficients for FA-1 firing were significantly greater than zero for all three stimulus parameters and contributed to median IFR similarly (Fig. 8A), while SA-1 firing was reliably determined by force and, to a lesser extent, velocity (Fig. 8B).

**Figure 8.**
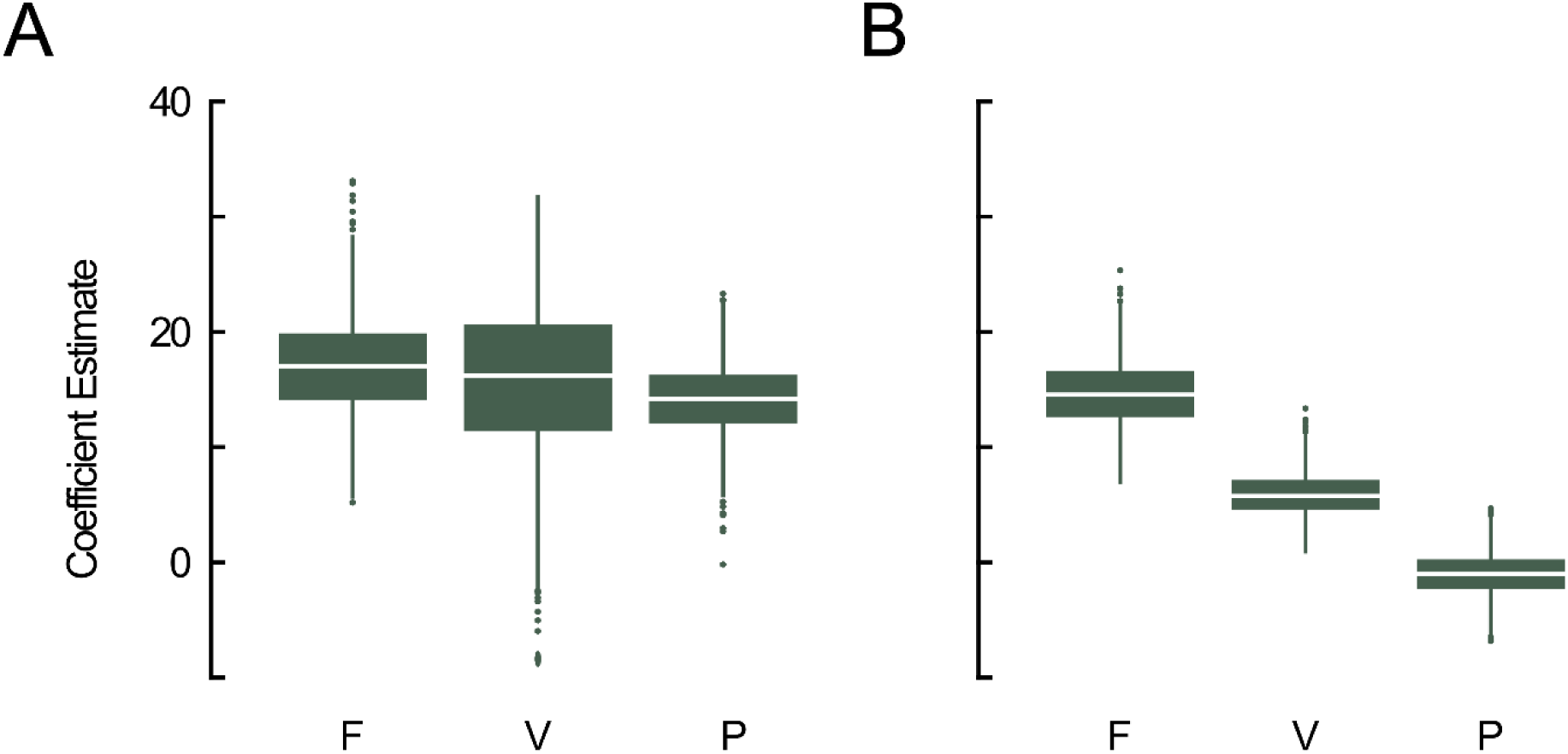
Bootstrapped coefficient estimates show that stimulus parameters with significant contribution to the median IFR are not centered around zero. That is, (A) FA-1 median IFR is determined by force, velocity, and spatial period, whereas (B) SA-1 median IFR is determined by force and velocity. The lower and upper edges of the box demarcate the interquartile range, and the white line indicates the median. Whiskers delimit the minimum and maximum values before the inner and outer fences (not illustrated).

## Discussion

A longstanding view in tactile neuroscience is that each class of LTM afferent is specialized for one dominant stimulus feature, for example, SA-1 afferents act as pressure sensors, or FA-2 detect vibration. Our results challenge this reductionist perspective by showing that individual LTMs on the finger are not exclusively dedicated to single stimulus features, but instead display overlapping response properties. Such overlap suggests that tactile information is represented across afferent populations in a distributed manner, where each class contributes differently-weighted inputs to completely capture the details of the stimulus. This multidimensional encoding framework helps explain how the human hand achieves its dexterity and precision in object interaction.

Investigations of LTM afferent responses in glabrous hand skin have established their fundamental properties^8,15,20,22–26^, but fewer studies have modeled how single human afferents encode dynamic stimuli. We take this further by comparing three different parameters directly in FA-1 and SA-1 mechanoreceptive afferents. Foundational work by Johnson and colleagues demonstrated that afferent firing patterns convey rich information about vibration, motion, and textured surfaces through both firing rate and precise temporal spiking^11,12,25,27,28^. Subsequent studies further explored the spatiotemporal receptive fields and population coding of tactile stimuli, highlighting the role of precise spike timing in supporting fine texture and motion perception^14,18,29–33^. These studies established the importance of dynamic encoding, but largely focused on single stimulus dimensions, where this is clearly more complicated in everyday tactile interactions. While the four classes of LTM afferents in glabrous skin have distinct receptive field structures and temporal response properties, functional overlap likely exists: when one unit fails to reliably encode a stimulus feature, another may compensate. In this vein, our results provide new human data showing that FA-1 afferents reliably integrate information about normal force, sliding velocity, and spatial period, underscoring their function as dynamic event detectors and slip encoders^22,34^, and their critical role. However, it further suggests that these mechanoreceptors function as general-purpose encoders of dynamic tactile events, supporting the role of FA-1 afferents to provide a stream of continuous input during touch, helping to determine grip force during object slip and giving motion cues during active, real-time object manipulation and surface exploration.

We showed that SA-1 afferents primarily encoded the magnitude of normal force and responded modestly to sliding velocity, consistent with their established role as detectors of static or slowly-changing skin deformations, and supports their function in fine force discrimination and pressure detection. In classic primate studies on tactile spatial encoding, SA-1 afferents were shown to produce spatially-stable firing patterns that encoded the geometry of raised-dot and grating stimuli, suggesting a role in encoding fine spatial detail and edges^10,11^. However, our results indicate SA-1 dynamic sensitivity to variations in force and velocity. Philips et al. ^11^ showed that, SA-1 firing increased markedly with higher contact force and the onset of movement, despite their lower dynamic sensitivity relative to FA-1 afferents. This indicates that SA-1 afferents are not purely static encoders of spatial form, but are influenced by the interaction between skin and stimulus. In our experiments, the absence of localized indentations—as with periodic gratings instead of raised dots—suggests that SA-1 firing primarily reflects the distributed strain field or local pressure variations rather than discrete edge features. This interpretation aligns with a recent finding emphasizing that tactile afferents encode local skin deformation and strain rather than geometric features^35^. Thus, the well-established association of SA-1 afferents with edge detection in the monkey may stem from their high sensitivity to localized indentation forces, which correlate with edges in many classic stimuli, but are not intrinsic to the receptor’s encoding mechanism, and edge-related firing should not be taken as a defining or sufficient characteristic of SA-1 afferents. Their responses emerge from general sensitivity to local strain and pressure, meaning that many forms of contact—beyond geometric edges—can evoke comparable activity provided the stimulus produces adequate skin deformation. Differences in SA-1 afferent density and skin mechanics between monkeys and humans may further contribute to variations in spatial resolution and dynamic response profiles, possibly explaining why our SA-1 firing rates (∼40 Hz) are substantially lower than those observed in primate dot-stimulus experiments (∼120 Hz^12^). Together, FA-1 and SA-1 afferents therefore provide complementary information about both the intensity and temporal structure of tactile stimuli during sliding contact.

During the present experiments, care was taken to stimulate the center of each afferent’s receptive field, but it is unlikely that every unit experienced identical mechanical stimulation. Variations in receptive field geometry, microneurography electrode placement, and local skin mechanics inevitably introduce subtle differences across recordings. Nonetheless, the observed variability in firing rates across trials and units likely very much reflects intrinsic differences in afferent tuning rather than purely methodological artifacts. This variability contributes to the broader symphony of tactile encoding: even within a single afferent class, units are not homogeneous. As revealed by our models, individual FA-1 and SA-1 units exhibit diverse sensitivities to normal force, sliding velocity, and spatial period. While the overall patterns align with canonical descriptions of each class, the richer picture from our regression analyses highlights that afferents distribute their coding capacity across multiple stimulus dimensions. This functional heterogeneity supports the view that the precise, real-time tactile representation during object manipulation arises from the coordinated activity of partially overlapping, yet individually distinct, afferents—a coordinated ensemble whose collective firing encodes the nuanced features of textured surfaces.

Despite the utility of our descriptive statistics and models, limitations should be noted. Microneurography remains labor-intensive and yields sparse datasets, due to the challenge of isolating and maintaining stable recordings from individual fibers. This was especially true for the present study, where the desired location of the receptive fields was the center of the fingertip, and longer recordings were essential, allowing multiple repetitions of different combinations of stimulus parameters. Afferent types cannot be preselected for during the nerve search process, and the lower density of SA-2 and FA-2 afferents in glabrous skin is reflected in our limited sampling, which precluded meaningful modeling of these types. Type 2 LTMs make up around a third of mechanoreceptors on the hand. However, on the glabrous fingertip, the density of type 1 units is much higher, meaning that, in reality, around 90% of mechanoreceptive afferents found here are type 1^8^. Therefore, although this highlights the need for future work to expand knowledge about type 2 LTM afferent responses, it is clearly the type 1 LTMs that are contributing most to tactile encoding and perception at the fingertip. To mitigate some data sparsity, bootstrapping provided a validation of the robustness of the FA-1 and SA-1 mixed-effects models by empirically estimating sampling variability through resampling. However, bootstrap estimates inherit the limitations of the original dataset: namely, its size and the distribution of recorded units. Extending these models with additional recordings remains crucial to capture the coordinated activity underpinning tactile discrimination, as does adding further conditions outside our tested ranges.

The multilevel models developed here provide a quantitative framework for predicting single-unit firing rates under varying stimulus conditions. These numerical illustrations may inform tactile encoding schemes for biomimetic sensors or neural prostheses designed to restore naturalistic touch feedback. For example, by incorporating realistic predictions of how afferent types respond to dynamic touch, future prostheses could emulate the nuanced tactile cues required for secure grip and dexterous manipulation, as well as provide precise signals about texture. It is evident from our work that touch at the fingertip entails the mimicry of the type 1 LTM afferents, therefore these should be focused on in neuroprosthetic applications, taking into account their complex firing to different touch parameters. Furthermore, our models lay the groundwork for exploring how populations of afferents jointly encode touch. In everyday interactions, hundreds, if not thousands, of afferent units are activated simultaneously, and it is the symphony of firing that produces the rich picture of tactile percepts, including the incorporation of temperature that ‘colors’ the sensation. A practical follow-up would explore the ranges in which each afferent type most effectively encodes stimulus information, and in the liminal spaces, determine the functional overlap between afferent types. Future experiments could examine the role of temperature on mechanoreceptive afferent firing during tactile interactions, as well as exploring direct recordings from thermoreceptors, although these are notoriously challenging to record from.

Interactions with ordinary objects presents a multitude of textural features over many orders of magnitude. The synthesis of these features constitutes the extraordinary human ability to discriminate minute textural differences and effortlessly manipulate objects in hand. Providing intuitive tactile feedback in hand prostheses has the potential to improve their functionality and acceptance^36,37^, by restoring touch and reducing reliance on vision during object manipulation. By advancing our understanding of how human tactile afferents respond to multi-parameter dynamic stimuli, our findings demonstrate that single human LTM afferents systematically encode multiple aspects of dynamic surface contact, underscoring their essential roles in real-time tactile discrimination. The statistics presented in this work contribute to the knowledge base needed to design more naturalistic haptic interfaces and prosthetic feedback systems. Ultimately, mapping the single-afferent coding strategies brings us closer to replicating the remarkable precision and adaptability of the human sense of touch.

## Materials and Methods

### Participants

The study was approved by the local ethics committee at the University of Gothenburg (368-06, 628-17) and the experiments were conducted in accordance with the Declaration of Helsinki, apart from pre-registration. The participants received verbal and written information prior to the experiment and gave written informed consent. Data were collected from a total of 18 healthy human volunteers (14 female, 4 male), 22-32 years old (mean 24.7 years).

### Set-up and nerve recordings

Participants were seated comfortably in a reclining dental chair with their left arm naturally extended and resting with support, with the ventral side down. For increased stability, the arm was fixated with a vacuum pillow. The setup allowed for access of the upper arm for peripheral nerve recordings, as well as access of the distal halves of the index, middle, and ring fingers, on the volar side of the hand, for tactile stimulation using a mechatronic platform.

Human axonal in vivo microneurography recordings from single LTM afferents in the left median nerve were captured using high-impedance, insulated tungsten needle electrodes (FHC Inc., Bowdoin, ME, USA; 50 mm length, 0.2 mm shaft diameter). After insertion of the electrode, the experimenter incrementally modified the electrode’s position and lightly stroked the participant’s hand until an intraneural location was reached and single unit impulses from an LTM afferent could be discerned from background activity. Units were classified as one of four types: FA-1, FA-2, SA-1, or SA-2, according to their response to indentation and the size and existence of well-defined borders of their receptive fields^9^, i.e., slowly-adapting afferents continue to fire during sustained pressure, whereas fast-adapting do not, and type 1 afferents have small, point-like receptive fields, whereas type 2 are much larger and less defined. SA-1 and SA-2 units were further distinguished by their responses to a long-lasting (30 s) 10 mN indentation with a von Frey monofilament at the center of the receptive field, where SA-2 units exhibit a more regular discharge pattern compared to SA-1^38^. The nerve signal was recorded using a bandpass filter set to 0.2-4.0 kHz and sampled at 12.8 kHz using Zoom/SC (University of Umeå, Umeå, Sweden). Additional details about the nerve search procedure can be found in^39^.

### Protocol

Controlled mechanical stimulation of the identified unit was executed using a custom-built mechatronic platform, controlled via a LabVIEW (National Instruments, Austin, TX, USA) program designed to deliver normal forces of 100, 200, 400, or 800 mN over 20 mm distance at velocities of 5, 10, 20, or 40 mm/s. The robotic platform was used to slide mounted periodic gratings under these exact conditions across the receptive field of the identified afferent unit on the finger, which was immobilized and secured by its nail, glued to a holder, to ensure precise stimulation. Additional details about the robotic platform can be found in^40^.

The periodic gratings, sized 35 x 32 mm, were manufactured in TUFSET rigid polyurethane plastic with varied spatial periods defined as the distance between the leading edges (Unilever R&D, Port Sunlight, UK). For “fine” gratings with spatial periods between 280 – 520 μm, each ridge had a width and depth of 100 and 300 μm, respectively. For “coarse” gratings with spatial periods between 1280 – 1920 μm, each ridge had a width and depth of 400 and 1200 μm, respectively. Gratings tested had 17 different spatial periods: 280, 390, 400, 410, 420, 440, 480, 520, 1280, 1440, 1520, 1560, 1600, 1640, 1680, 1860, and 1920 μm.

Two gratings with different spatial periods were glued adjacent to one another to a metal plate, which was then fixed to the robotic platform, in order to expedite testing (see Figure 1). One side of the plate could be tested, then the other side. The platform disengaged one grating before contacting the second, preventing overlap in recordings. Each grating was unidirectionally passed across the receptive field of the afferent unit for a minimum of 6 repetitions. Pairs of gratings on the plates were changed between conditions, to test other spatial periods, and the order of stimuli was pseudo-randomized. The complete set of parameters yielded 272 unique combinations of stimulus settings, although not all conditions could be tested for every unit recorded due to the limited (and essentially random) time that a stable recording from the unit could be achieved.

### Analysis

#### Microneurography signal pre-processing

The detection of single action potentials in the recorded microneurography signal was performed via in-house-developed scripts implemented in MATLAB (The MathWorks, Inc., Natick, Massachusetts, USA). The script classifies the peak of nerve spikes based on local minima and maxima within a 1.2 ms window^41^ using an optimal linear filter^42^. The automated spike detection was verified by the experimenter to remove artifacts and to reject data where the signal appears to constitute multi-channel inputs. The resultant point process, derived from the peaks of the nerve spikes, served as the input sequence for the analysis to follow.

#### Statistics and data modeling

Descriptive statistics for the mean, median, and peak instantaneous firing rate (IFR) and the coefficient of variation were computed in MATLAB. Firing rates were not normalized in order to preserve unit-specific differences. Median IFR curves were generated and fit using non-linear least squares regression.

Regression analyses were conducted in R Statistical Software (v4.5.0; R Core Team 2025). To address heteroscedasticity, predictor variables—normal force (F), sliding velocity (V), and spatial period (P)—were log-transformed. This transformation linearizes potentially multiplicative relationships by converting multiplicative effects on the original scale to additive effects on the log scale. This facilitates interpretation of coefficients as elasticities (i.e., percentage change in outcome per percentage change in predictor)^43,44^. This also helps meets the regression assumption of approximately constant residual variance, which is often violated in biological data^45^. The outcome variable (median IFR) was left untransformed.

To quantify the joint effect of stimulus parameters on median IFR, separate linear regressions were fit for each unit. For each unit *j*, the model was specified as:

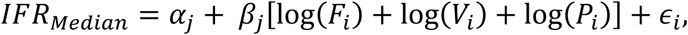

where *i* indexes trials within unit *j*; α is the regression intercept; β is the regression coefficient, and ϵ is the residual error term. In R notation: *IFR*_*median*_ ∼ log(*F*) + log(*V*) + log(*P*).

To estimate the overall effect of each predictor across units, we fitted linear mixed-effects models using the *lme4* package (v.1.1-37)^46^. Mixed-effects models account for repeated measures and within-unit correlations by including random intercepts, improving generalizability across the sample^44^. The following 7 nested models were compared to evaluate the contribution of each predictor:

1. Models with one predictor: log(*F*); log(*V*); log(*P*),
2. Models with pairs of predictors: log(*F*) + log(*V*); log(*F*) + log(*P*); log(*V*) + log (*P*),
3. Full model with all predictors: log(*F*) + log(*V*) + log (*P*).

The full mixed-effects model was specified as:

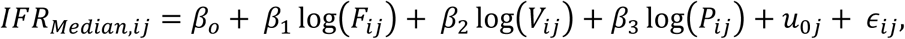

where *i* are the trials within each unit *j*; β_0_ is the fixed intercept; β_1_, β_2_, β_3_ are fixed slopes; *u*_0*j*_ is the random intercept for unit *j*, assumed to follow *N*(0, σ_u_^2^); and ϵ *ij* ∼ *N*(0, σ^2^) is the residual error. In R notation: *IFR*_*median*_ ∼ log(*F*) + log(*V*) + log(*P*) + (1 | *Unit ID*).

Model performance was evaluated using the Akaike Information Criterion (AIC) and marginal *R*^2^, using the *MuMin* package (v1.48.11). The AIC balances model fit and complexity by penalizing the inclusion of additional parameters; lower AIC values indicate a model that better explains the data without overfitting^47^. Models were ranked by AIC, and the relative AIC differences from the best-fitting model were reported. Fixed-effect estimates and 90% confidence intervals were visualized to illustrate the estimated influence of each stimulus parameter.

To test robustness of fixed-effect estimates and capture sampling variability, we performed non-parametric bootstrapping with 1000 resamples using the *boot* package (v.1.3-31)^48^. Each resample was drawn with replacement, the model refitted, and new fixed-effect estimates computed. This approach, which does not assume normally distributed residuals, provides empirical distributions of parameter estimates, particularly useful with limited data^49,50^. Boxplots were used to summarize the spread and central tendency of these bootstrapped estimates, where the median, as is indicated by a middle line, and marker the lower and upper edges of the box demarcate the interquartile range.

## Data Availability

Due to the sensitive nature of human participant data, the dataset generated and analyzed during the current study is not publicly available but can be accessed upon reasonable request.

## Acknowledgments

The study was funded by grants to JW from the Swedish Research Council (K2007-63X-03548; K2010-62X-03548; 2021-02552) and from the European Commission (EU FP6 NANOBIOTACT 33287). VAL was funded by the EU Horizon 2020 Marie Sklodowska-Curie Action Programme NEUTOUCH No 813713. We would like to thank Karin Göthner for excellent technical support.

